# Lithium-induced ciliary lengthening sparks Arp2/3 complex-dependent endocytosis

**DOI:** 10.1101/2022.04.18.488674

**Authors:** Brae M Bigge, Prachee Avasthi

## Abstract

Ciliary length is highly regulated across cell types, but this tight regulation can be disrupted by lithium, which causes ciliary elongation across cell types and organisms. Here, we use the powerful ciliary model *Chlamydomonas reinhardtii* to investigate the mechanism behind lithium-induced ciliary elongation. Protein synthesis is not required for lengthening, and the target of lithium is GSK3, which has substrates that can influence membrane dynamics. Further, in addition to elongation of the microtubule core, ciliary assembly requires a supply of ciliary membrane. To test if the membrane for ciliary lengthening could be from the Golgi or the cell body plasma membrane, we treated cells with either Brefeldin A or Dynasore respectively. Cilia were able to elongate normally with Brefeldin treatment, but Dynasore treatment resulted in defective lengthening. Genetic or acute chemical perturbation of the Arp2/3 complex, which is required for endocytosis in these cells, blocks lithium-induces ciliary lengthening. Finally, we looked at filamentous actin in lithium-treated cells and found an increase in Arp2/3 complex-and endocytosis-dependent puncta near the base of cilia. Blocking endocytosis by inhibiting the Arp2/3 complex or dynamin, confirmed by visual loss of endocytic structures, prevents lithium-induced ciliary elongation. We previously reported that endocytosis was required for early ciliary assembly from zero length, and here, we demonstrate that endocytosis is also required for ciliary elongation from steady state length. Thus, we hypothesize that lithium-induced ciliary elongation occurs through a mechanism that involves a supply of additional ciliary membrane through endocytosis.

## INTRODCUTION

The plasma membrane-ensheathed, microtubule-based cilium is important for signaling and motility, and defects in this organelle can lead to a large number of diseases termed ciliopathies (Reiter and Leroux 2017). Thus, ciliary length is tightly regulated across cell types. However, the mechanisms by which such regulation occurs are still being investigated. Many studies have focused on perturbations that result in shorter cilia, but the relatively infrequent examples of ciliary lengthening can hold the keys to uncovering regulatory mechanisms.

An example of a perturbation that increases ciliary length is lithium. The ciliary elongation elicited by lithium treatment is ubiquitous across cell types, occurring in *Chlamydomonas* (Nakamura, Takino, and Kojima 1987; Wilson and Lefebvre 2004), chondrocytes (Soave et al. 2022; Thompson et al. 2016), ependymal cells (Kong et al. 2015), human fibroblast-like synoviocytes (Ou et al. 2009), fibroblasts (Ou et al. 2012), mouse brains (Miyoshi et al. 2009), NIH3T3 cells (Miyoshi et al. 2009), human induced pluripotent stem cell-derived neurons (Miki et al. 2019), Sertoli cells from pig testes (Ou et al. 2014), and others. Lithium is thought to target GSK3 (Wilson and Lefebvre 2004), but the mechanism by which this causes ciliary elongation are relatively unknown.

Much of what we know about ciliary length regulation comes from studies in the unicellular green algae *Chlamydomonas reinhardtii*. *Chlamydomonas* serves as a powerful model for studying length regulation and assembly of cilia as it has two persistent and symmetric cilia that are structurally and mechanistically similar to the cilia of mammalian cells. The cilia of *Chlamydomonas* can be easily severed and regrown in a matter of hours which allows for dissection of the complex processes involved in ciliary assembly and regulation (Jack and Avasthi 2018; Rosenbaum, Moulder, and Ringo 1969; Paul A. Lefebvre 1995; P. A. Lefebvre et al. 1978). Further, genetic mutants for *Chlamydomonas* exist for 83% of the nuclear genome (Li et al. 2019; Cheng et al. 2017). These mutants can be used to identify important genes required for ciliary length regulation. For example, short flagella mutants (JARVIK et al. 1984, 1) and long flagella mutants (Barsel, Wexler, and Lefebvre 1988; Asleson and Lefebvre 1998; Nguyen, Tam, and Lefebvre 2005, 1; L.-W. Tam, Wilson, and Lefebvre 2007) have been used to help researchers better understand the mechanisms involved in maintaining ciliary length.

Using *Chlamydomonas* several models for ciliary length regulation have been proposed (Ludington et al. 2015; Avasthi and Marshall 2012; Ishikawa and Marshall 2017a; Marshall 2015), including the limiting-precursor model (Rosenbaum, Moulder, and Ringo 1969), a few diffusion-based models (Levy 1974; Ludington et al. 2015; Hendel, Thomson, and Marshall 2018), the molecular ruler model (Marshall 2015), the time-of-flight model (Ishikawa and Marshall 2017b), the mechanosensitive ion channel model (Besschetnova et al. 2010; Beck and Uhl 1994), the swim speed feedback model (D. Tam and Hosoi 2011; Osterman and Vilfan 2011), and the balance point model (Marshall and Rosenbaum 2001; Marshall et al. 2005). The bulk of what we know regarding ciliary length regulation is related to the availability or turnover of protein, generally tubulin, but cilia are not merely composed of proteins. They are also ensheathed in plasma membrane. This leads to questions about whether membrane could be limiting. In fact, when cells are treated with Brefeldin A to collapse the Golgi and prevent the delivery of Golgi-derived membrane, cilia shorten (Dentler 2013). Further, data suggest that the Arp2/3 complex and actin are involved in reclamation of membrane from the cell body plasma membrane that is required for normal ciliary assembly (Bigge et al. 2020).

Because lithium causes ciliary lengthening across a broad range of cells and organisms, we use it as a powerful tool to investigate the mechanisms that result in ciliary elongation in *Chlamydomonas* and beyond. We investigate the trafficking mechanisms that deliver the excess membrane required for additional growth past steady state length and find that endocytosis is important for ciliary elongation induced by lithium. Further, using both chemical inhibitors and genetic mutants, we find a role for Arp2/3 complex-mediated actin networks in ciliary lengthening induced by lithium. Altogether, we propose a new model where the additional membrane required for lithium-induced ciliary lengthening is endocytosed in an Arp2/3 complex-dependent manner.

## RESULTS

### Inhibition of GSK3 by many mechanisms results in ciliary elongation

It has been proposed that lithium targets GSK3 in *Chlamydomonas* (Wilson and Lefebvre 2004). We questioned whether inhibition of GSK3 was directly responsible for ciliary elongation seen with lithium. To answer this, we turned to other GSK3 inhibitors. Three classes of GSK3 inhibitors exist: metal cations that interfere with ATP binding, ATP completive inhibitors, and non-ATP competitive inhibitors. Lithium is thought to inhibit GSK3 by competing with magnesium ions required for ATP binding and by phosphorylation. To confirm that GSK3 inhibition is the cause of ciliary elongation caused by lithium, we employed inhibitors from each of the other two classes. We used CHIR99021 and (2’Z,3’E)-6-Bromoindirubin-3’-oxime (6-BIO) as ATP competitive inhibitors and Tideglusib as a non-ATP competitive inhibitor.

Consistent with previous results, treatment with LiCl induces ciliary elongation through either interference with ATP binding or through phosphorylation of GSK3 (**Figure 1A**). The ATP competitive inhibitor CHIR99021 has been shown to modestly elongate cilia in foreskin fibroblasts (Ou et al. 2012). We confirmed that in *Chlamydomonas* 100 μM CHIR99021 also results in an increase in ciliary length (**Figure 1B**). Next, we looked at the competitive inhibitor 6-BiO. Although 2 μM 6-BiO has been used in *Chlamydomonas* and caused ciliary shortening (Kong et al. 2015), when we treated cells with 100 nM 6-BIO, we observed ciliary elongation (**Figure 1C**). Finally, we looked a non-competitive inhibitor of GSK3, Tideglusib, for which the effects on cilia have not been previously observed. When cells were treated with 20 μM of Tideglusib, we saw an increase in ciliary length consistent with other methods of GSK3 inhibition (**Figure 1D**). While each inhibitor may have its own set of of-targets, they all share a unique on-target of GSK3, lending further support to GSK3 being the cilium length relevant target of GSK3. This also suggests that the method of inhibition of GSK3 is not important for ciliary lengthening. Whether GSK3 was inhibited via competition for ATP binding or phosphorylation, cilia were able to elongate.

**Figure 1.**
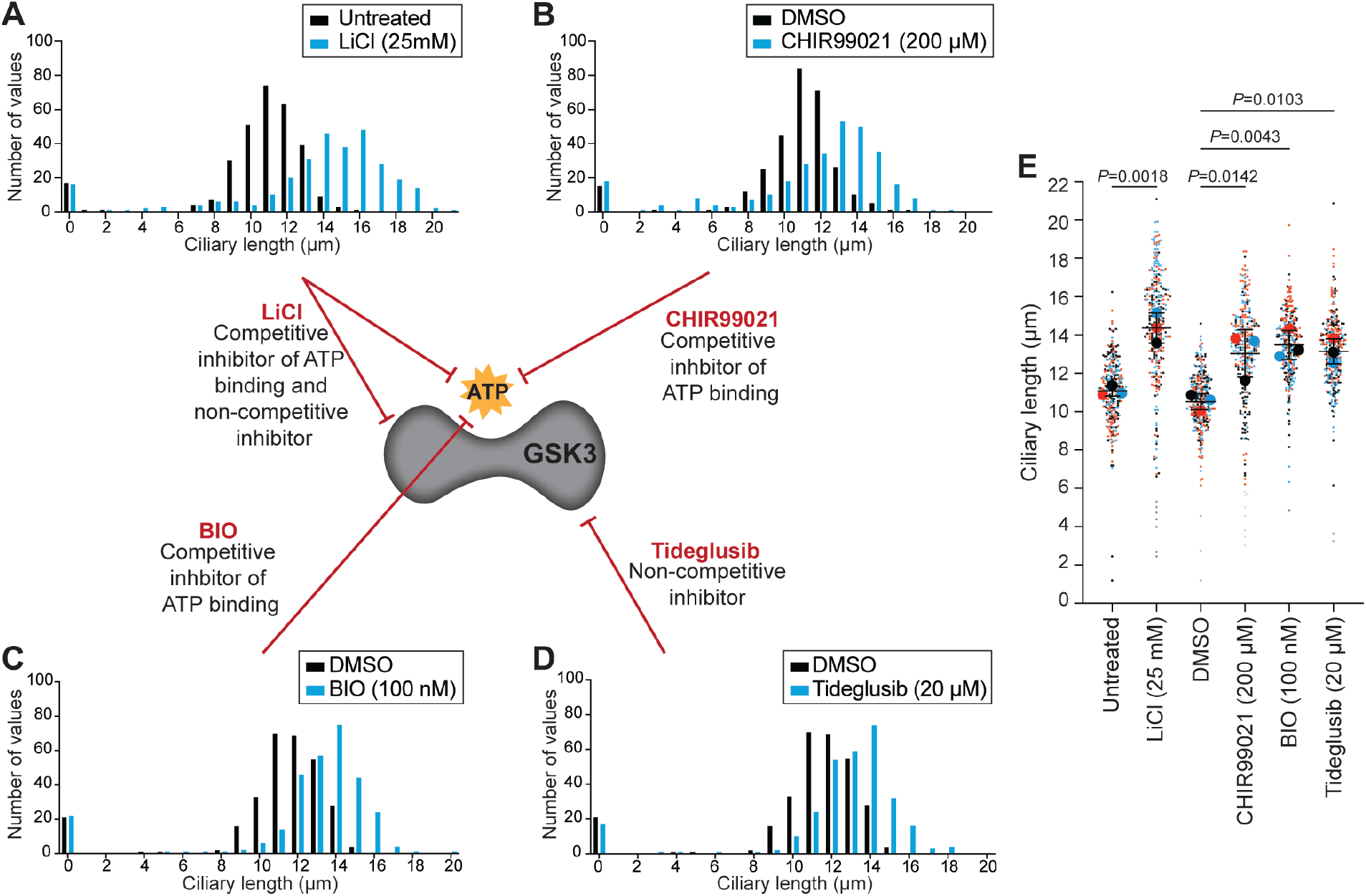
Inhibition of GSK3 through multiple mechanisms results in ciliary elongation. A-D) Wild-type (CC-5325) cells were treated with GSK3 inhibitors including 25 mM LiCl (A), 200 μM CHIR99021 (B), 20 μM Tideglusib (C), or 100 nM BIO (D) for 90 minutes. Histograms show distribution of ciliary lengths. n=100 cells in 3 separate biological replicates. **E)** Dot plot representing the data in A-D excluding zero length cilia. Each color represents a separate biological replicate where n=100. The large dots represent means from each replicate and the mean (solid lines), standard deviation (error bars), and statistical analyses were calculated using those means. P values shown on the plot are the results of a one-way ANOVA followed by Tukey’s multiple comparisons analysis.

### Membrane for lithium-induced ciliary elongation comes from endocytosis

To better understand the mechanism behind ciliary elongation induced by GSK3 inhibition, we looked at the source of the ciliary material. The cilium is primarily composed of the microtubule-based axoneme and the membrane surrounding the organelle. Thus, for the rapid growth seen with lithium, the cell needs to deliver not only protein, but also membrane to the cilia during a relatively short time frame. Treatment of cells with lithium and cycloheximide, which blocks protein synthesis, resulted in normally elongating cilia, suggesting that new protein synthesis is not required for lithium-induced ciliary elongation and the protein required for assembly must come from a pool somewhere in the cell (Wilson and Lefebvre 2004) (**Supplemental Figure 1**).

The membrane required for ciliary elongation induced by lithium must come from one of two sources: the Golgi, which is generally thought to be the main source of ciliary membrane (Nachury, Seeley, and Jin 2010; Rohatgi and Snell 2010), or a pool in the cell body plasma membrane (Bigge et al. 2020). To differentiate between these two possibilities, we treated cells with either Brefeldin A (BFA), a drug that causes Golgi collapse, or Dynasore, a drug that interferes with dynamin-mediated endocytosis. When cells were treated with BFA to block membrane delivery form the Golgi, cilia were still able to elongate normally in LiCl (**Figure 2A-B**). This suggests that membrane from the Golgi is not required for lithium-induced elongation.

**Figure 2.**
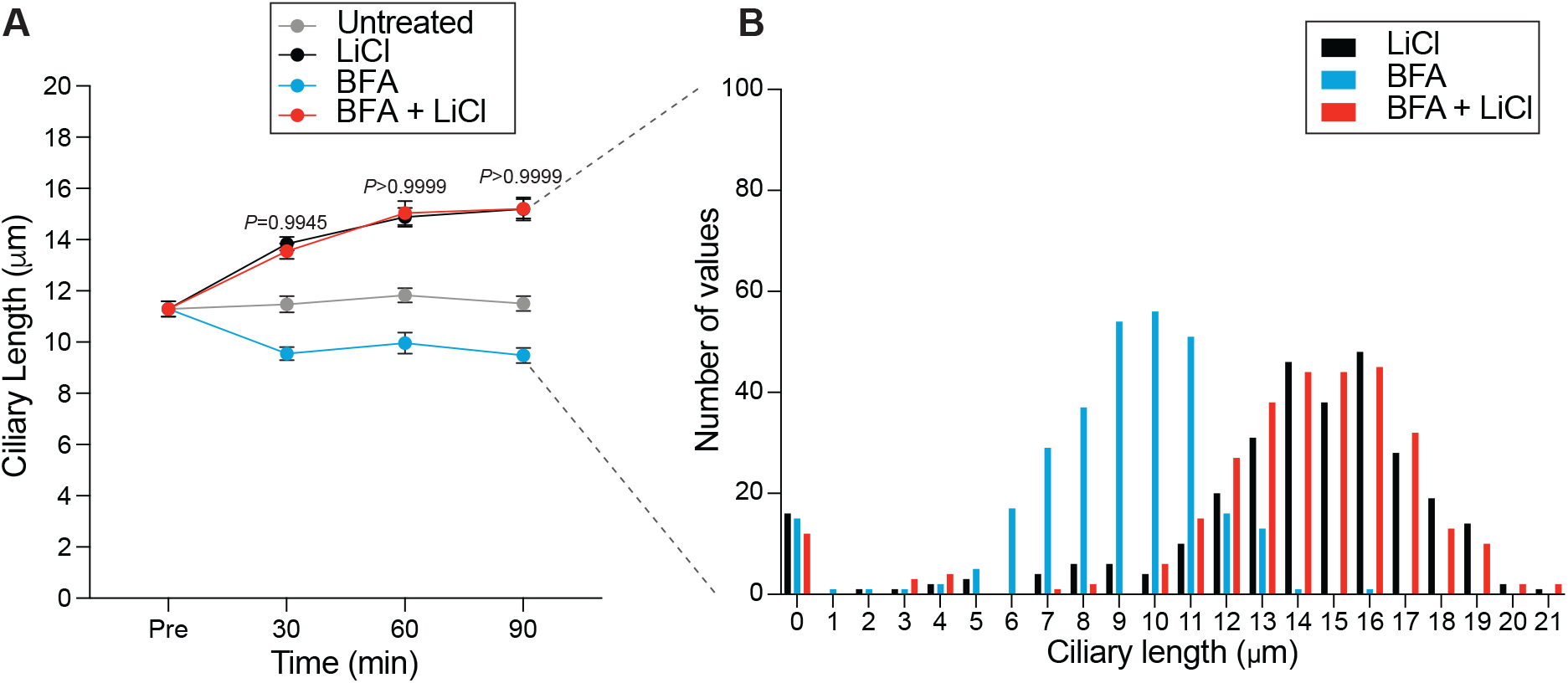
**Golgi-derived membrane is not required for lithium-induced ciliary elongation**. A)Wild-type (CC-5325) cells were treated with 36 μM Brefeldin A or 36 μM Brefeldin A with 25 mM LiCl. n=30 cells per time point and sample in 3 separate biological replicates. Significance was determined using a two-way ANOVA with a Tukey’s multiple comparisons test. Values above the lines show the comparison between LiCl and BFA with LiCl. Each time point is ns at the p-values listed on the graph. Additionally, cells treated with BFA alone have significantly (p<0.0001) shorter cilia than untreated cells, confirming function of BFA. **B)** The 90-minute time point from (A) was expanded to n=100 cells for each biological replicate. The histograms show distribution of ciliary lengths.

Conversely, when treated with Dynasore to inhibit endocytosis, ciliary elongation was defective (**Figure 3A-B**), implying endocytosis is required for lithium-induced elongation and that endocytosis requires dynamin. To further probe the involvement of endocytosis in ciliary elongation, we turned to the dynamin family. GSK has been previously shown to target the dynamin protein family; in neuronal and non-neuronal mammalian cells dynamin 1 is usually inactive due to phosphorylation by GSK3ß, but when cells are treated with CHIR99021 to inhibit GSK3, dynamin 1 is dephosphorylated and endocytosis rates increase significantly (Srinivasan et al. 2018; Smillie and Cousin 2012). Meanwhile, GSK3α has been found to phosphorylate mammalian Dynamin 2 (Laiman et al. 2021).

**Figure 3.**
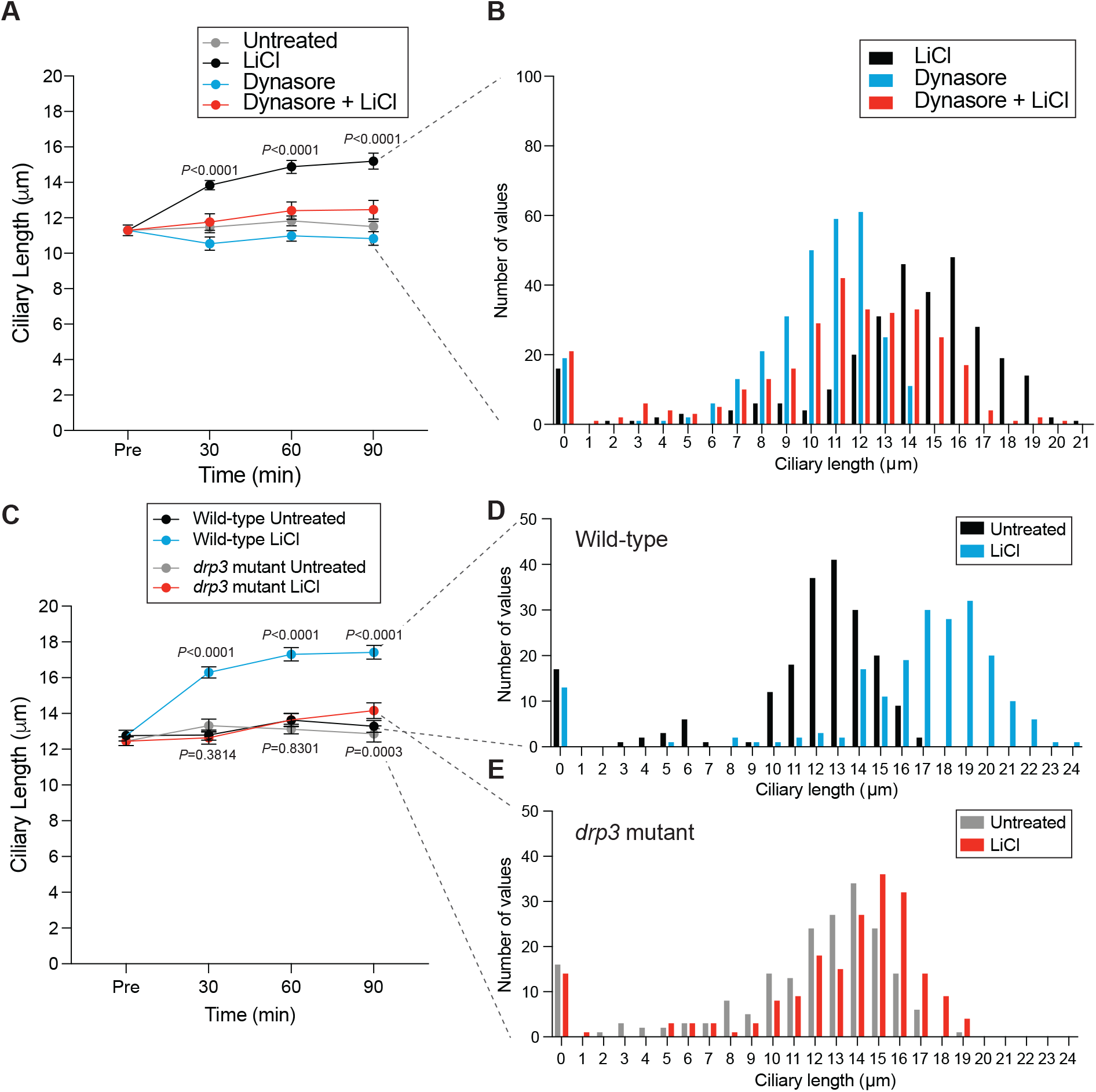
Lithium-induced ciliary elongation requires endocytosis. A)Wild-type (CC-5325) cells were treated with 100 μM Dynasore or 100 μM Dynasore with 25 mM LiCl. n=30 cells per time point and sample in 3 separate biological replicates. Significance was determined by two-way ANOVA with a Tukey’s multiple comparisons test. P values on the graph are comparing LiCl treated cells to cells treated with LiCl and Dynasore. Each time point is significantly different (****). **B)** The 90-minute time point from (A) was expanded to n=100 cells for each biological replicate. The histograms show distribution of ciliary lengths. **C)** Wild-type (CC-5325) cells and *drp3* mutant cells were treated with 25 mM LiCl. n=30 cells per time point and sample in 2 separate biological replicates. Significance was determined by two-way ANOVA with a Tukey’s multiple comparisons test. P values on the graph are comparing Wild-type cells treated with LiCl to *drp3* mutant cells treated with LiCl. Each time point is significantly different (****). Additionally, the P values below the line compare untreated *drp3* mutant cells with LiCl-treated *drp*3 mutant cells. The 30- and 60-minute time point are not significantly different, by the 90-minute time point is (***)**. D-E)** The 90-minute time points from (D) were expanded to n=100 cells for each biological replicate. (D) represents wild-type cells and (E) represents *drp*3 mutant cells. The histograms show distribution of ciliary lengths.

Canonical dynamins have not been identified in *Chlamydomonas*, but the genome contains many dynamin related proteins (DRPs). DRPs differ from dynamins in that they do not contain all 5 traditional dynamin domains: A GTPase domain, a middle domain, a pleckstrin homology domain, a guanine exchange domain, and a proline rich domain (Elde et al. 2005) (**Supplemental Figure 2**). *Chlamydomonas* contains 9 DRPs with similarity to a canonical dynamin (DRP1-9). The DRP with the highest similarity to canonical dynamin is DRP3 (**Supplemental Figure 2**). To determine if GSK3 could be a potential kinase for this protein, we employed ScanSite4.0, which confirmed that of the 9 DRPs of *Chlamydomonas*, the only one with a traditional GSK3 target sequence was DRPs (**Supplemental Figure 2**).

To investigate whether DRP3 might be involved in the ciliary elongation that results from lithium treatment, we obtained a mutant from the *Chlamydomonas* mutant library (CLiP) (Cheng et al. 2017; Li et al. 2019). This mutant has a cassette inserted early in the gene. When we treated these cells with lithium, ciliary elongation was decreased (**Figure 3C-D**), suggesting that this DRP3 is involved in the ciliary response to GSK3 inhibition. Elongation was not fully blocked by this mutation in DRP3, this is potentially due to the presence of other DRPs that might compensate for the lost function of DRP3.

### Lithium-induced ciliary elongation promotes formation of filamentous actin puncta

The high demand for new membrane and protein for ciliary elongation led us to hypothesize that treatment with GSK3 inhibitors would lead to an increase in actin dynamics. Previously, we found that upon deciliation and rapid initial ciliary assembly, filamentous actin puncta visualized with phalloidin and dependent on the Arp2/3 complex form at the apex of the cell near the cell body plasma membrane (Bigge et al. 2020). These puncta are reminiscent of endocytic pits seen in yeast and require the Arp2/3 complex, which is known to be involved in endocytosis in cells with cell walls, like yeast and *Chlamydomonas* (Bigge et al. 2020; Basu, Munteanu, and Chang 2014; Aghamohammadzadeh and Ayscough 2009; Carlsson and Bayly 2014). Thus, we phalloidin stained cells to visualize filamentous actin and these punctate structures in cells treated with GSK3 inhibitors. Untreated cells formed some puncta at the apex of the cell (**Figure 4A**), but treatment with either lithium (LiCl), CHIR99021, 6-BIO, or Tideglusib increased the percentage of cells with dots and the number of dots per cell (**Figure 4A-B**). *arpc4* mutant cells never form dots confirming that the formation of these actin puncta is Arp2/3-dependent (**Figure 4A**). The increased formation of these Arp2/3 complex-dependent filamentous actin puncta suggests a burst of endocytosis triggered by inhibition of GSK3 occurring during times or rapid ciliary elongation.

**Figure 4.**
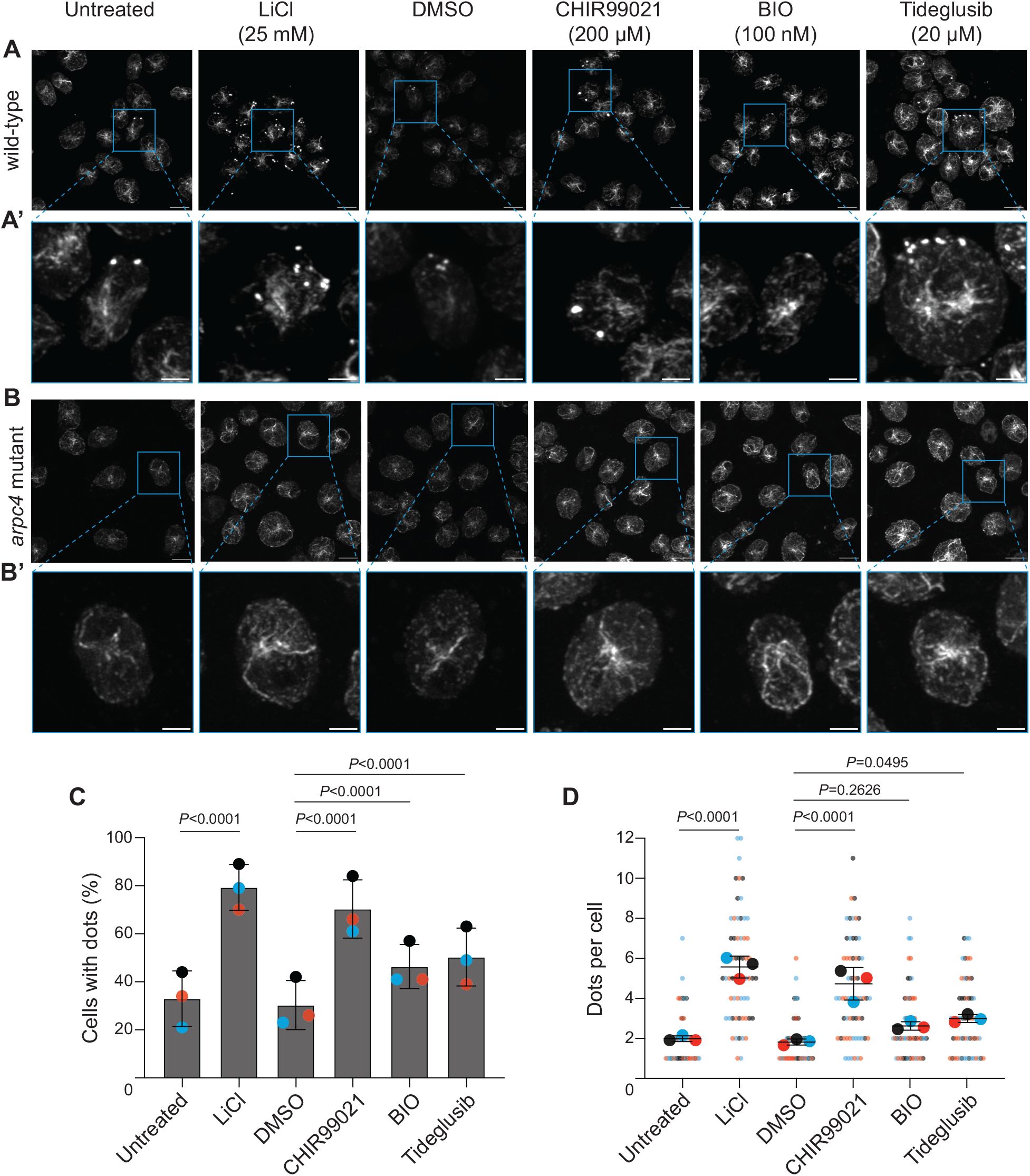
GSK3 inhibition increases Arp2/3 complex-dependent filamentous actin puncta. A)Maximum intensity projections of z stacks taken of wild-type cells treated with either nothing, LiCl (25 mM), DMSO, CHIR99021 (200 μM), BIO (100 μM), or Tideglusib (20 μM) and stained with phalloidin to visualize filamentous actin. Scale bars represent 5 μm. **A’)** Zoomed in images of the boxed cells in A. **B)** Maximum intensity projections of z stacks taken of *arpc4* mutant cells treated with either nothing, LiCl (25 mM), DMSO, CHIR99021 (200 μM), BIO (100 μM), or Tideglusib (20 μM) and stained with phalloidin to visualize filamentous actin. Scale bars represent 5 μm. **B’)** Zoomed in images of the boxed cells in B. **C)** Quantification of the percentage of cells with dots in either untreated, lithium treated, DMSO treated, CHIR99021 treated, BIO treated, or Tideglusib treated samples. n=100 in 3 separate biological replicates. Significance was determined using Chi Square analysis and Fisher’s exact tests. **D)** Quantification of the number of dots per cell in cells treated with either DMSO, CHIR99021, BIO, or Tideglusib at the specified concentrations. n=20 cell in 3 separate biological replicates. Significance was determined using a one-way ANOVA and a Tukey’s multiple comparisons test.

To further investigate the increase in actin puncta, we employed an mNeonGreen-tagged Lifeact peptide, which labels filamentous actin populations in live cells. Using this method, we were able to visualize dynamic actin and puncta in both untreated and lithium-treated wild-type cells (**Figure 5A, Supplemental Videos 1-2**). Then using the ImageJ/FIJI Plugin Trackmate, we tracked the movement of filamentous actin accumulations within the cell (Ershov et al. 2021; Tinevez et al. 2017). We found that actin dynamics were significantly increased in wild-type cells treated with lithium compared with untreated cells (**Figure 5B-C**). The increased actin dynamics in lithium-treated cells were particularly enriched near the membrane.

**Figure 5.**
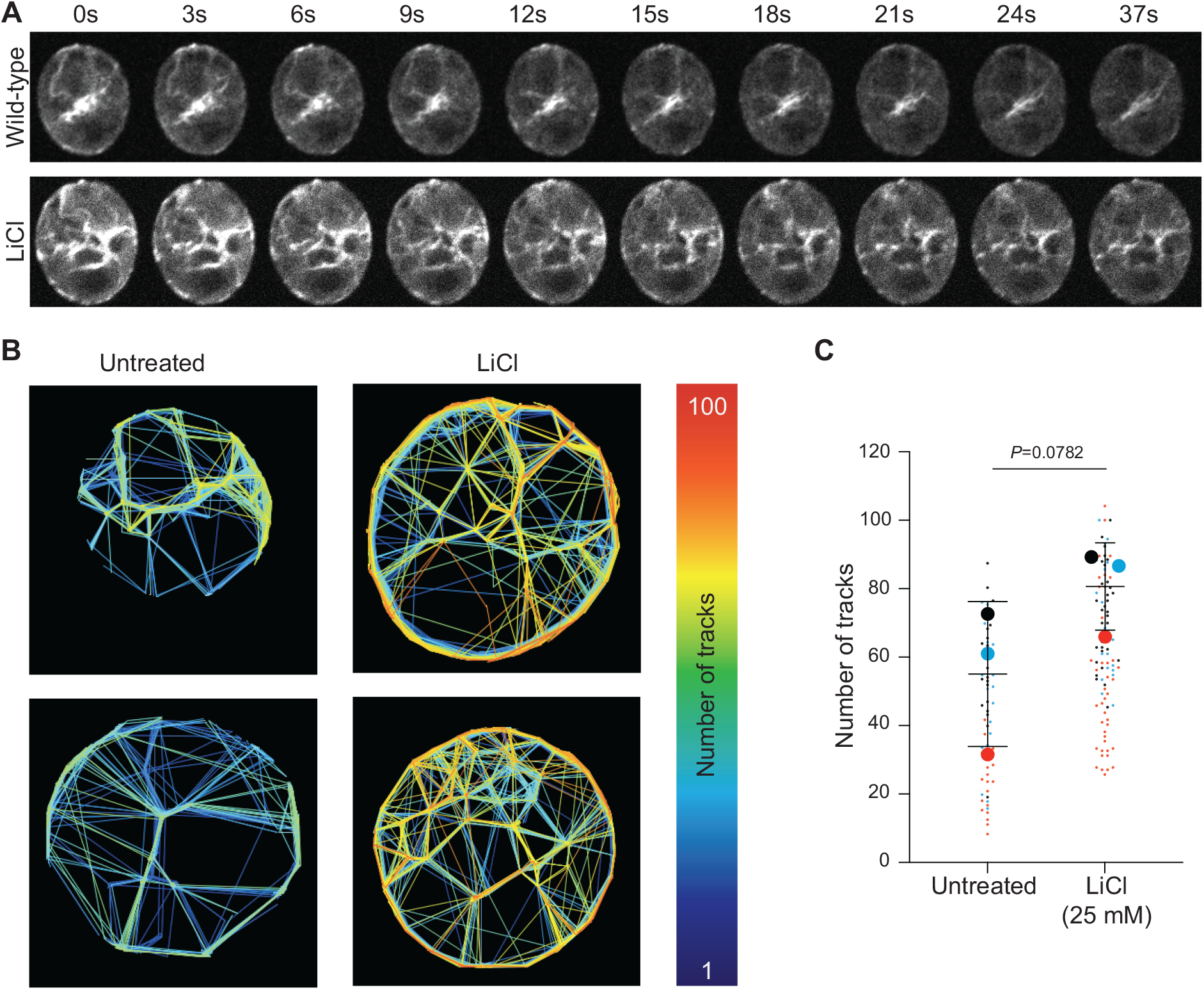
Actin dynamics increase near the membrane during lithium-induced ciliary elongation. A)Time-lapse representation of videos taken of cells expressing Lifeact-mNeonGreen either untreated or treated with 25 mM LiCl and then immediately imaged on a Nikon Spinning Disk microscope for 30 seconds. **B)** Videos represented in (A) were analyzed using the FIJI plugin, TrackMate. This identifies spots in videos and tracks their movement allowing us to measure actin dynamics. The tracks created in TrackMate were adjusted so that cells with less tracks will have more blue tracks, but cells with more tracks will start to have tracks with more oranges and reds according to the scale. **C)** From the TrackMate data, we found the number of tracks in untreated cells and cells treated with LiCl. Each color represents one of 3 separate biological replicates. The large dots represent the means from each biological replicate with the line and error bars showing the mean and standard deviation of the means from each replicate. Two-tailed t-test was performed on the means to give the p-value on the graph.

### The Arp2/3 complex is required for lithium-induced ciliary elongation

The increase in Arp2/3 complex-dependent actin puncta with lithium or GSK3 inhibition led us to question whether the Arp2/3 complex was required for ciliary elongation induced by lithium. Further, we previously showed that the Arp2/3 complex is required for rapid ciliary assembly in the initial stages of ciliogenesis (Bigge et al. 2020), leading us to hypothesize that the Arp2/3 complex will also be required for rapid ciliary elongation induced by lithium. Chemical inhibition of the Arp2/3 complex with the small molecule inhibitor CK-666 resulted in defective ciliary elongation in lithium (**Figure 6A-B**). We confirmed this result with genetic inhibition of the Arp2/3 complex component ARPC4 using an *arpc4* mutant first described in Bigge et al. 2020 (**Figure 6C-E**). This suggests the Arp2/3 complex is required for lithium-induced ciliary elongation.

**Figure 6.**
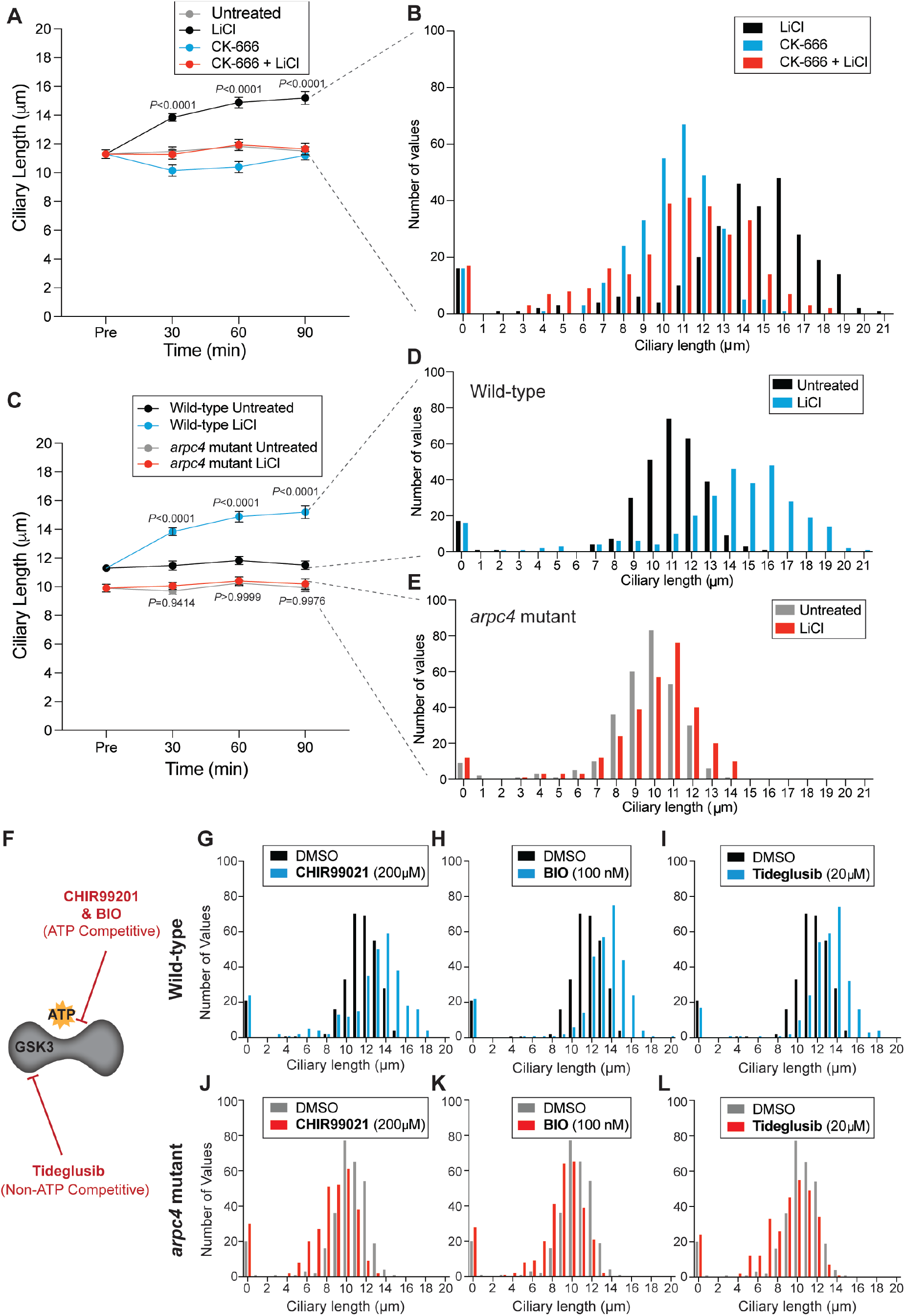
The Arp2/3 complex is required for ciliary elongation induced by GSK3 inhibition. A)Wild-type (CC-5325) cells were treated with 100 μM CK-666 or 100 μM CK-666 with 25 mM LiCl. n=30 cells per time point and sample in 3 separate biological replicates. Significance was determined by one-way ANOVA and a Tukey’s multiple comparisons test. The p values above the lines show the comparison between cells treated with LiCl and cells treated with LiCl and CK-666. **B)** The 90-minute time point from (A) was expanded to n=100 cells for each biological replicate. The histograms show distribution of ciliary lengths. **C)** Wild-type (CC-5325) cells and *arpc4* mutant cells were treated with 25 mM LiCl. n=30 cells per time point and sample in 3 separate biological replicates. Significance was determined by one-way ANOVA and a Tukey’s multiple comparisons test. The p values above the lines show the comparison between wild-type cells treated with LiCl and *arpc4* mutant cells treated with LiCl. The p values below the lines show the comparison between untreated *arpc4* mutant cells and *arpc4* mutant cells treated with LiCl. At each time point, this comparison was not significant. **D-E)** The 90-minute time points from (C) were expanded to n=100 cells for each biological replicate. (D) represents wild-type cells and (E) represents *arpc4* mutant cells. The histograms show distribution of ciliary lengths. **F)** Schematic showing GSK3 inhibitory functions of the 3 additional inhibitors. BIO and CHIR99021 are ATP competitive while Tideglusib is ATP non-competitive. **G-I)** Wild-type cells were treated with either 200μM CHIR99021 (B), 100nM BIO (C), or 20μM Tideglusib (E) for 90 minutes. Histograms show distribution of ciliary lengths. n=100 cells in 3 separate biological replicates. **J-L)** *arpc4* mutant cells were treated with either 200μM CHIR99021 (E), 100nM BIO (F), or 20μM Tideglusib (G) for 90 minutes. Histograms show distribution of ciliary lengths. n=100 cells in 3 separate biological replicates.

To confirm that this requirement of the Arp2/3 complex is connected to the inhibition of GSK3, we also treated wild-type and *arpc4* mutant cells with CHIR99021, BIO, and Tideglusib. In all cases, wild-type cells were able to elongate while *arpc4* mutant cells either did not elongate or shortened (**Figure 6G-L**). Thus, the mechanism whereby GSK3 inhibition results in ciliary elongation requires the Arp2/3 complex.

## DISCUSSION

In this work, we investigate the mechanisms behind ciliary elongation induced by lithium treatment in an effort to better understand the factors regulating ciliary length. Cilia are primarily composed of a microtubule axoneme and a plasma membrane that contains the axoneme. Many of the studies to understand ciliary length regulation focus on the microtubule axoneme, the amount of free tubulin available for assembly, and the delivery of tubulin to and from the ciliary tip through intraflagellar transport (IFT). While these are important factors and we are interested in how our data can fit with these existing models, we focus instead on the plasma membrane that ensheathes the axoneme. Because new protein synthesis is not required for ciliary elongation induced by lithium (Wilson and Lefebvre 2004) (**Supplemental Figure 2**), we hypothesized that the primary source of membrane for elongation was perhaps separate from the Golgi, which is typically thought to be the primary source of membrane for ciliary assembly. Previous work has shown that while the Golgi is required for ciliary maintenance and assembly (Dentler 2013), it is not the only source of membrane. Instead, membrane reclaimed through actin and Arp2/3-complex dependent endocytosis are required for ciliary assembly or growth from zero length (Bigge et al. 2020). This led us to question whether that same mechanism might be required for ciliary elongation from steady state length induced by lithium treatment.

We found that GSK3 inhibition resulted in increased ciliary length using CHIR99021, BIO, Tideglusib, and Lithium which each target GSK3 through different mechanisms (**Figure 1**). CHIR99021, BIO, and Tideglusib were designed for use in humans, but their ability to elongate cilia in *Chlamydomonas* suggests that they are able to target GSK3 ubiquitously and that this results in ciliary elongation across organisms. Next, we showed that while Golgi-derived membrane was not required for ciliary lengthening, endocytosis and dynamin function were needed to elongate cilia (**Figures 2–3**). We could not however rule out other sources of membrane, such as the endosomal network. Finally, we showed that GSK inhibition resulted in a burst of Arp2/3-complex dependent dots and increased actin dynamics at the membrane (**Figure 4–5**). These actin dots were previously observed during initial ciliary assembly when there is a high demand for membrane that can be quickly incorporated into cilia (Bigge et al. 2020). They are reminiscent of endocytic patches or pits seen in yeast where actin is required for endocytosis to overcome turgor pressure related to the presence of a cell wall, which *Chlamydomonas* also has (Aghamohammadzadeh and Ayscough 2009; Basu, Munteanu, and Chang 2014; Carlsson and Bayly 2014) While we cannot say these are *Chlamydomonas* endocytic pits with the current data, we do believe they speak to the presence of actin functioning at the membrane during these periods when membrane is in high demand. Finally, we show that the Arp2/3 complex is absolutely required for lithium-induced ciliary lengthening from steady state (**Figure 6**), again suggesting an actin- and Arp2/3 complex-dependent process is required for ciliary elongation. We hypothesize that this Arp2/3 complex-dependent process is linked to endocytosis, but direct endocytosis during lithium treatment has not been observed in these cells.

Based on our data, we propose a model for lithium induced ciliary elongation were lithium targets GSK3 which results in a burst of Arp2/3 complex-dependent endocytosis to reclaim membrane for ciliary lengthening (**Figure 7**). While we made strides toward identifying the membrane-related targets of GSK3 that result in ciliary elongation, we do not suggest that this is the only pathway targeted by GSK3 that contributes to ciliary elongation. An interesting next step would be to further dissect the targets of GSK3 that also contribute to this elongation. For example, some mechanism must be at play that increases the recruitment of proteins that compose the axoneme and the IFT machinery. Additionally, it would be interesting to determine if these phenotypes observed in *Chlamydomonas* are conserved in other organisms that elongate their cilia in lithium.

**Figure 8.**
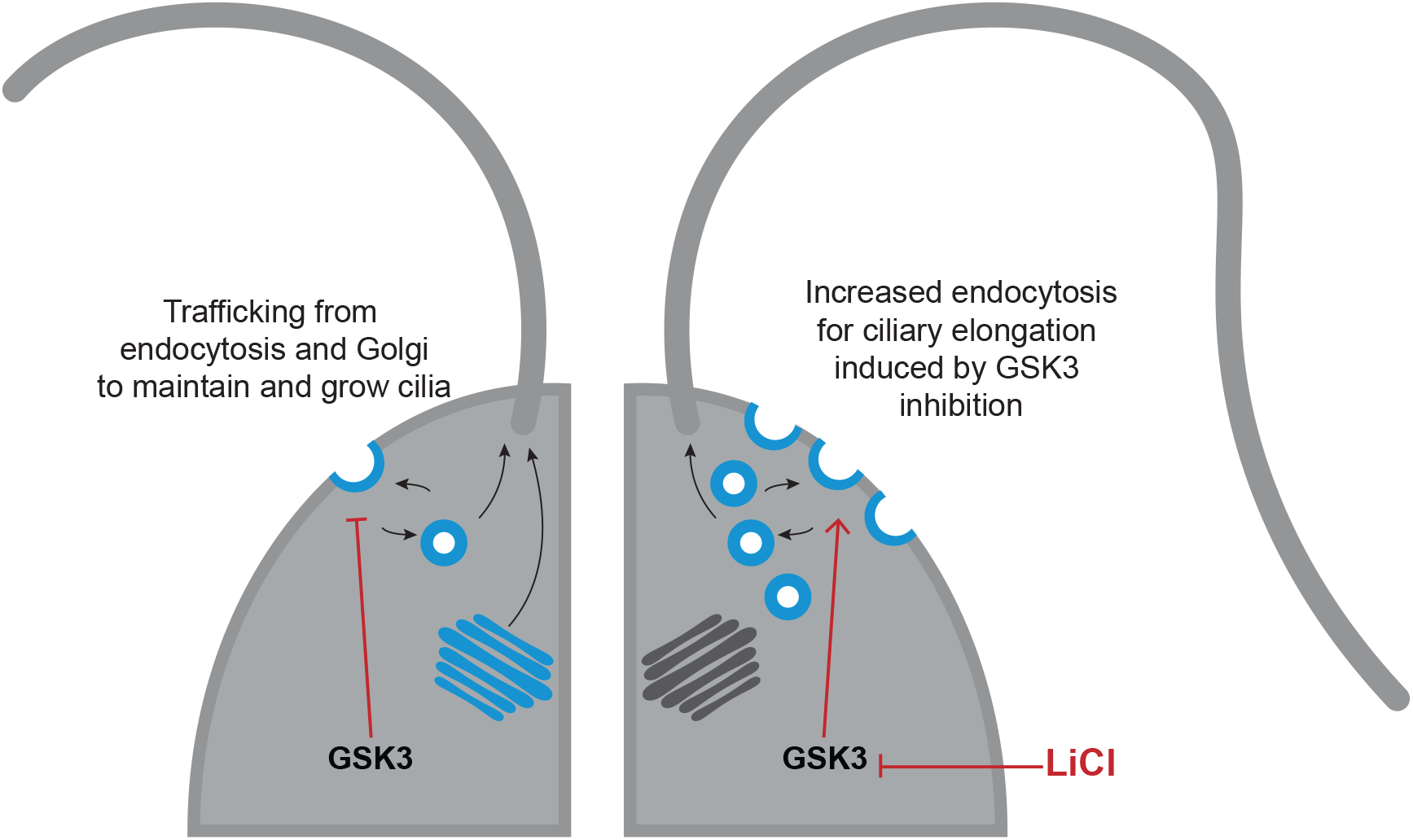
Lithium targets GSK3 which results in a burst of endocytosis to reclaim membrane for ciliary elongation. During normally ciliary maintenance and assembly, Golgi-derived membrane and some plasma membrane-derived membrane are transported to the cilium. However, during GSK3 inhibition, there is a burst of endocytic activity that is necessary to quickly grab membrane for ciliary elongation.

Finally, because these data are focused on the ciliary membrane instead of the ciliary axoneme, they provide new insight and lend support to the models of ciliary regulation that have already been established as follows (Ludington et al. 2015; Avasthi and Marshall 2012; Ishikawa and Marshall 2017a; Marshall 2015). The limiting-precursor model suggests that cells make exactly enough ciliary precursor proteins to form cilia of a certain length. However, even without new protein synthesis cilia are able to grow to half length (Rosenbaum, Moulder, and Ringo 1969), and in our data, cilia are able to immediately elongate their cilia well beyond steady-state length when treated with lithium. Therefore, in order for our lithium data and the regeneration in cycloheximide to fit the limiting pre-cursor model would require additional mechanisms to allow for growth past 1 cilium length like inhibition of autophagy, sequestration, and/or degradation during lithium treatment. Alternatively, it is possible that what is limiting cilium growth is not protein, but membrane, consistent with our data. The time-of-flight model proposes that there is a degradable signal, like phosphorylation, that is incorporated into IFT trains that move into the cilia. The longer the IFT trains with the signal are in the cilia, the more time for the signal to be degraded, thus providing a readout of ciliary length. However, no active length measurer has been identified (Ishikawa and Marshall 2017b). In lithium treated cells, where cilia are much longer than usual, the degradable signal would normally be fully degraded by the time it reached the base of the cilia, but for this model to hold, lithium treatment would have to interfere with the degradation of this signal. Another model suggests that mechanosensitive ion channels in the ciliary membrane regulate ciliary length because the longer the cilia is the more ion channels can be present in the membrane and the more ions can be imported into the cell (Besschetnova et al. 2010; Beck and Uhl 1994). This model is interesting to consider as lithium itself is an ion, Li^+^. It is therefore possible that the presence of lithium interferes with ion balance, such as that of Ca^2+^,to regulate cilium length (Besschetnova et al. 2010). The swim speed feedback model proposes that the cilia grow to an optimal length for swimming and that the length therefore depends on fluid forces upon the cilia (D. Tam and Hosoi 2011; Osterman and Vilfan 2011). Lithium treated cells are unable to swim as their cilia are not only elongated but also paralyzed (Wilson and Lefebvre 2004; Dentler 2005). This could affect the sensing of fluid forces in the cilia and cause feedback that could control ciliary length, although paralyzed flagella mutants do not display elongated cilia, suggesting that should this model be relevant lithium would have a different mechanism of paralysis. Our data could also support other models of ciliary length regulation that deal more with the axoneme if lithium is simultaneously affecting both the membrane and the axoneme. These models include the tubulin diffusion-based model (Levy 1974; Craft Van De Weghe et al. 2020), the signal diffusion-based model (Ludington et al. 2015), the kinesin diffusion-based model (Hendel, Thomson, and Marshall 2018), the molecular ruler model (Marshall 2015), and finally the balance point model, which considers the constant turnover of tubulin at ciliary tips and the fact that ciliary assembly is not a linear process, but instead slows as cilia lengthen. This model suggests that there is a point where the decreasing assembly rate and the constant disassembly rate where cilia reach a steady state length (Marshall and Rosenbaum 2001; Marshall et al. 2005).

Altogether, we show that GSK3 inhibition sparks Arp2/3 complex and actin-dependent endocytosis to reclaim membrane for ciliary elongation. These data fit well with the some of the proposed models for ciliary regulation outlined above. Some interesting next steps include determining other targets of GSK3 inhibition that might contribute to ciliary elongation, investigating whether this pathway is conserved across organisms as lithium induced ciliary elongation is, and uncovering new data that might help lithium-induced ciliary lengthening and our data fit with established and possibly new models of ciliary elongation.

## MATERIALS AND METHODS

### Strains

The *arpc4* mutant (LMJ.RY0402.232713), *drp3* mutant (LMJ.RY0402.215697), and the wild-type parent strain (CC-5325) are from the *Chlamydomonas* Resource Center. The *arpc4* mutant was confirmed previously (Bigge et al. 2020). The *drp3* mutant was confirmed using 2 primer pairs. The first pair included: AGAAGGCCAGTTTCTCCTCGG and TTAAGCTCGACCTCCCTCAA. The second pair included: ATAGCCCGCCAAATCAGTCC and ACAGCAACACTGGTACACGC. Cells were grown and maintained on 1.5% Tris-Acetate Phosphate (TAP) agar plates. Prior to experiments, liquid TAP cultures were inoculated and grown overnight under constant red and blue light with agitation.

### Ciliary studies

Cells were treated with drugs at specified concentrations: 25 mM LiCl (MP Biomedicals, 194010), 200 μM CHIR99021 (Sigma, SML1046), 100 μM (2’Z,3’E)-6-Bromoindirubin-3’-oxime (Sigma, B1686), 20 μM Tideglusib (Sigma, SML0339), 10 μg/mL Cycloheximide (Sigma, C1988), 100 μM CK-666 (Sigma, 182515), 36 μM Brefeldin A (Sigma, B7651), and/or 100 μM Dynasore (Sigma, D7693). Cells were incubated with agitation under constant light for the specified times (usually 30 min, 60 min, and 90 min). Samples were taken prior to the experiment (‘Pre’) and at each time point by diluting 50 μl of cells 1:1 with 2% glutaraldehyde (EMS, 16220) and incubating at 4°C to allow cells to sediment. Cells were then imaged using a Zeiss Axioscope 5 DIC microscope at 40X (0.75 numerical aperture) and Zeiss Zen 3.1 (blue edition) software. Cilia were then measured in ImageJ using the segmented line and fit spline functions. One cilium per cell was measured as the cilia or the same cell should be equivalent lengths.

### Phalloidin staining

Procedure was originally published in (Craig and Avasthi 2019). Cells were allowed to adhere to poly-lysine treated coverslips and then fixed with 4% paraformaldehyde in 1X HEPES. Cells were then permeabilized with 80% acetone followed by 100% acetone before being allowed to dry fully. Coverslips rehydrated with PBS were then incubated with Phalloidin-Atto 488 (Sigma, 49409) for 16 minutes before being washed a final time with PBS. Coverslips were dried and then mounted with Fluoromount-G. Images were acquired using a Nikon Eclipse Ti-E microscope with a Yokogawa, two-camera CSU0W1 spinning disk system with a 100X oil-immersion objective lens (1.45 numerical aperture). Z-stacks were obtained using Nikon Elements. Then, maximum intensity projections were created in ImageJ. The number of cells with dots and the number of dots per cell were manually counted.

### Live cell imaging

The plasmid containing the Lifeact peptide tagged with mNeonGreen, pMO654, was a generous gift from Masayuki Onishi and is detailed in (Onishi et al. 2019). The plasmid was transformed into CC-5325 cells using electrophoresis. Briefly, cells were grown to an OD730 of 0.3-0.4 in liquid TAP media, pelleted, washed twice in Max Efficiency Buffer (Thermo), and finally resuspended to a volume of 250 μl Max Efficiency Buffer. This was divided into 2. To each, 1 μg of linearized plasmid was added. Cells with plasmid were incubated for 5 minutes at 4°C. The cells were then transferred to 4mm electroporation cuvettes. Using a BioRad Gene Pulser XCell set to exponential decay at 500V, 50 μF, and 800 Ohms, cells were electroporated. Following electroporation, cells were incubated at room temperature for 15 minutes then resuspended in 7 mL of liquid TAP + 40 mM sucrose and incubated overnight in the dark with constant agitation. The following day, cells were plated on 1.5% TAP plates with the appropriate selection antibiotic (Paromomycin). Colonies were selected, grown up, and tested by visually looking for fluorescence.

For imaging, the same Nikon Spinning Disk microscope described above was used. Cells, either untreated or immediately following lithium treatment, were imaged every 100 ms for 1 minute to create time series. The time series were then analyzed in ImageJ/FIJI using the TrackMate plugin (Tinevez et al. 2017; Ershov et al. 2021). The plugin identifies spots and then tracks their movement throughout the video. Only the first 300 frames of each image were used to eliminate concerns of bleaching or cell movement. Estimated object diameter was set to 1 μm and the quality threshold was set to 0.5. To make the colors of the track indicate how many tracks were present in each cell, they were colored by track ID with the highest track number value being set to the maximum value and the lowest track number value being set to the minimum.

## ACKNOWLEDGEMENTS

We want to express our appreciation to Masayuki Onishi for generously providing the Lifeact-mNeonGreen plasmid and the Avasthi lab for help throughout the project. We also want to thank the BioMT Core at Dartmouth College (NIH/NIGMS COBRE award P20-GM113132), the Genomics and Molecular Biology Shared Resources Core (NCI Cancer Center Support Grant 5P30CA023108-37), and Ann Lavanway for her help with microscopy. We also thank the Chlamydomonas Mutant Library Group at Princeton University, the Carnegie Institution for Science, and the Chlamydomonas Resource Center at the University of Minnesota for providing the indexed Chlamydomonas insertional mutant(s).

Finally, we thank our funding source, the NIGMS MIRA (R35GM128702).

## SUPPLEMENTAL MATERIAL

**Supplemental Figure 1.**
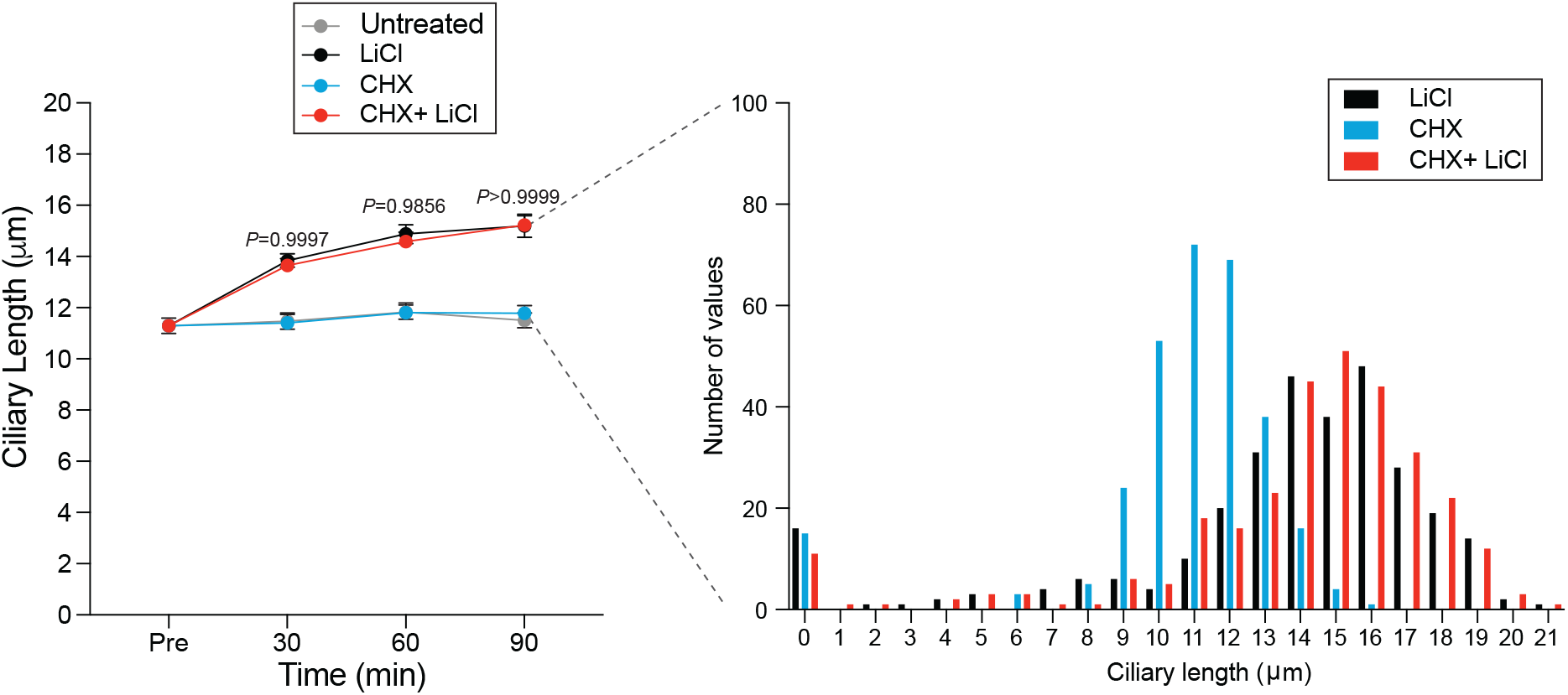
New protein synthesis is not required for lithium-induced ciliary elongation. **A)** Wild-type cells were treated with either 25 mM LiCl, 10 μM Cycloheximide (CHX), or a combination of the two drugs. n=30 for each sample at each time point in 3 separate biological replicates. Significance was determined by one-way ANOVA and a Tukey’s multiple comparisons test. The p values above the lines show the comparison between cells treated with LiCl alone or LiCl and CHX. In all cases, the comparison is not significant. **B)** A histogram representation of the 90-minute time point from (A). For this, n=100 for each sample in 3 separate biological replicates (300 total points).

**Supplemental Figure 2.**
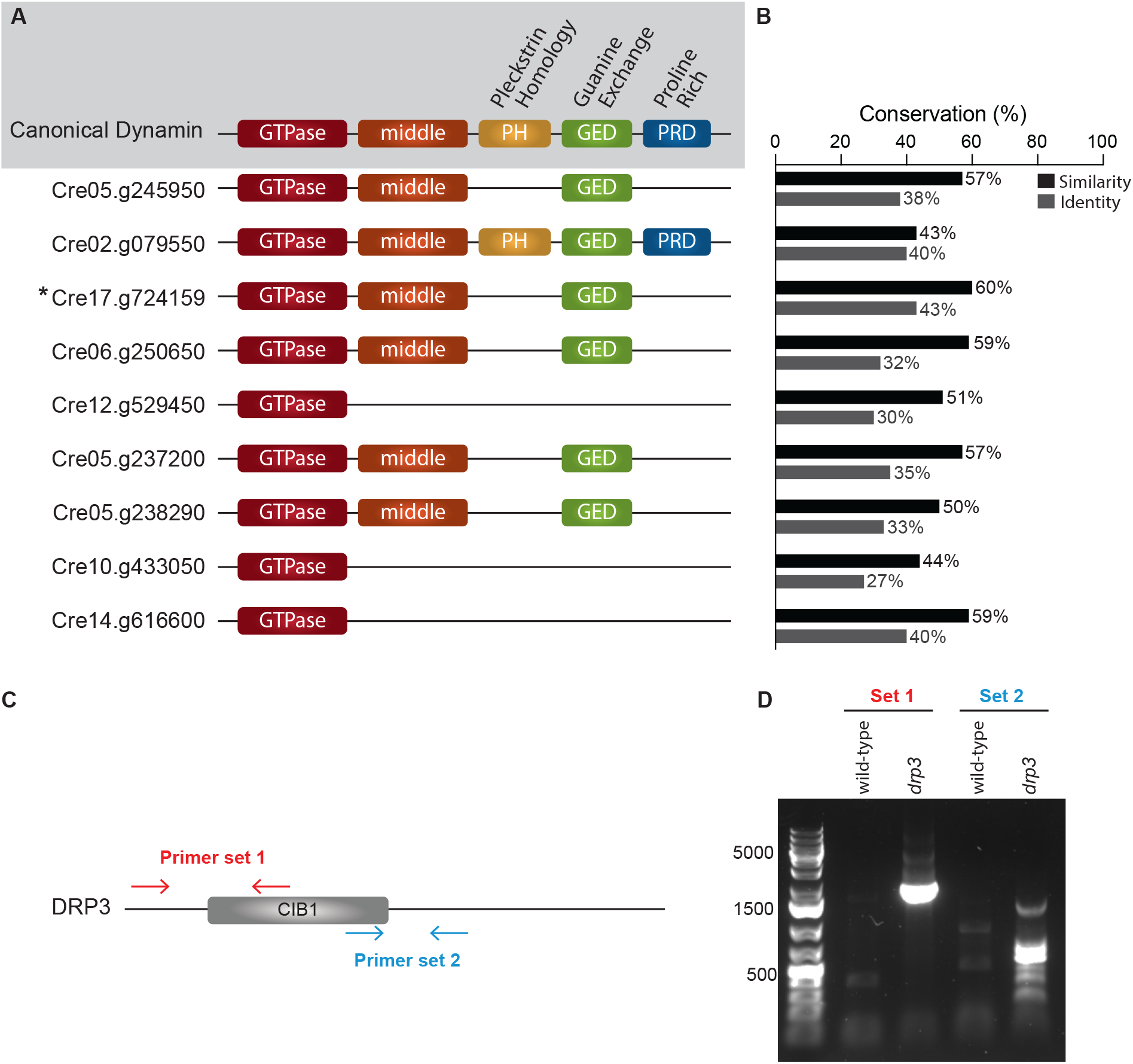
*Chlamydomonas* contains several dynamin related proteins (DRPs) with the most similar to canonical dynamin being DRP3 (Cre17.g724159) **A)** The canonical dynamin contains 5 domains represented in different colors above. The *Chlamydomonas* genome contains no conventional dynamins but does contain 9 dynamin related proteins (DRPs) that have various similarities compared with canonical dynamin. The *Chlamydomonas* DRPs above are represented by the gene identifier and are ordered by DRP number (the first is DRP1, the second is DRP2, etc). **B)** The graph on the right shows the similarity and identity of each DRP compared to canonical mammalian dynamin as determined by MUSCLE alignment. The DRP that is the most closely related to canonical dynamin is DRP3, Cre17.g724159. **C)** Schematic of the DRP3 genomic sequence with the primers used to confirm mutation represented by arrows. **D)** DNA gel electrophoresis showing resulting DNA sequences from the PCR using the primer sets shown in C.

## Notes

### Competing Interest Statement

P.A. is a paid consultant for Arcadia Science

## REFERENCES

Aghamohammadzadeh, Soheil, and Kathryn R. Ayscough. 2009. “Differential Requirements for Actin during Yeast and Mammalian Endocytosis.” Nature Cell Biology 11 (8): 1039–42. https://doi.org/10.1038/ncb1918.

Asleson, C. M., and P. A. Lefebvre. 1998. “Genetic Analysis of Flagellar Length Control in Chlamydomonas Reinhardtii: A New Long-Flagella Locus and Extragenic Suppressor Mutations.” Genetics 148 (2): 693–702. https://doi.org/10.1093/genetics/148.2.693.

Avasthi, Prachee, and Wallace F Marshall. 2012. “Stages of Ciliogenesis and Regulation of Ciliary Length.” Differentiation; Research in Biological Diversity 83 (2): S30–42. https://doi.org/10.1016/j.diff.2011.11.015.

Barsel, S. E., D. E. Wexler, and P. A. Lefebvre. 1988. “Genetic Analysis of Long-Flagella Mutants of Chlamydomonas Reinhardtii.” Genetics 118 (4): 637–48. https://doi.org/10.1093/genetics/118.4.637.

Basu, Roshni, Emilia Laura Munteanu, and Fred Chang. 2014. “Role of Turgor Pressure in Endocytosis in Fission Yeast.” Molecular Biology of the Cell 25 (5): 679–87. https://doi.org/10.1091/mbc.E13-10-0618.

Beck, Christina, and Rainer Uhl. 1994. “On the Localization of Voltage-Sensitive Calcium Channels in the Flagella of Chlamydomonas Reinhardtii.” The Journal of Cell Biology 125 (5): 1119–25.

Besschetnova, Tatiana Y, Elona Kolpakova-Hart, Yinghua Guan, Jing Zhou, Bjorn R Olsen, and Jagesh V Shah. 2010. “Identification of Signaling Pathways Regulating Primary Cilium Length and Flow-Mediated Adaptation.” Current Biology 20 (2): 182–87.

Bigge, Brae M, Nicholas E Rosenthal, David Sept, Courtney M Schroeder, and Prachee Avasthi. 2020. “Initial Ciliary Assembly in <em>Chlamydomonas</Em> Requires Arp2/3-Dependent Recruitment from a Ciliary Protein Reservoir in the Plasma Membrane.” BioRxiv, January, 2020.11.24.396002. https://doi.org/10.1101/2020.11.24.396002.

Carlsson, Anders E., and Philip V. Bayly. 2014. “Force Generation by Endocytic Actin Patches in Budding Yeast.” Biophysical Journal 106 (8): 1596–1606. https://doi.org/10.1016/j.bpj.2014.02.035.

Cheng, Xi, Gai Liu, Wenting Ke, Lijuan Zhao, Bo Lv, Xiaocui Ma, Nannan Xu, et al. 2017. “Building a Multipurpose Insertional Mutant Library for Forward and Reverse Genetics in Chlamydomonas.” Plant Methods 13 (1): 36. https://doi.org/10.1186/s13007-017-0183-5.

Craft Van De Weghe, Julie, J. Aaron Harris, Tomohiro Kubo, George B. Witman, and Karl F. Lechtreck. 2020. “Diffusion Rather than Intraflagellar Transport Likely Provides Most of the Tubulin Required for Axonemal Assembly in Chlamydomonas.” Journal of Cell Science 133 (17). https://doi.org/10.1242/jcs.249805.

Craig, Evan W., and Prachee Avasthi. 2019. “Visualizing Filamentous Actin Using Phalloidin in Chlamydomonas Reinhardtii.” Bio-Protocol 9 (12). https://doi.org/10.21769/BioProtoc.3274.

Dentler, William. 2005. “Intraflagellar Transport (IFT) during Assembly and Disassembly of Chlamydomonas Flagella.” The Journal of Cell Biology 170 (4): 649–59. https://doi.org/10.1083/jcb.200412021.

Dentler, William. 2013. “A Role for the Membrane in Regulating Chlamydomonas Flagellar Length.” PLOS ONE 8 (1): e53366. https://doi.org/10.1371/journal.pone.0053366.

Elde, Nels C, Garry Morgan, Mark Winey, Linda Sperling, and Aaron P Turkewitz. 2005. “Elucidation of Clathrin-Mediated Endocytosis in Tetrahymena Reveals an Evolutionarily Convergent Recruitment of Dynamin.” PLOS Genetics 1 (5): e52. https://doi.org/10.1371/journal.pgen.0010052.

Ershov, Dmitry, Minh-Son Phan, Joanna W. Pylvänäinen, Stéphane U. Rigaud, Laure Le Blanc, Arthur Charles-Orszag, James R. W. Conway, et al. 2021. “Bringing TrackMate into the Era of Machine-Learning and Deep-Learning.” BioRxiv, January, 2021.09.03.458852. https://doi.org/10.1101/2021.09.03.458852.

Hendel, Nathan L., Matthew Thomson, and Wallace F. Marshall. 2018. “Diffusion as a Ruler: Modeling Kinesin Diffusion as a Length Sensor for Intraflagellar Transport.” Biophysical Journal 114 (3): 663–74. https://doi.org/10.1016/j.bpj.2017.11.3784.

Ishikawa, Hiroaki, and Wallace F. Marshall. 2017a. “Intraflagellar Transport and Ciliary Dynamics.” Cold Spring Harbor Perspectives in Biology 9 (3). https://doi.org/10.1101/cshperspect.a021998.

Ishikawa, Hiroaki, and Wallace F. Marshall. 2017b. “Testing the Time-of-Flight Model for Flagellar Length Sensing.” Molecular Biology of the Cell 28 (23): 3447–56. https://doi.org/10.1091/mbc.e17-06-0384.

Jack, Brittany, and Prachee Avasthi. 2018. “Erratum to: Chemical Screening for Flagella-Associated Phenotypes in Chlamydomonas Reinhardtii.” Methods in Molecular Biology (Clifton, N.J.) 1795: E1. https://doi.org/10.1007/978-1-4939-7874-8_19.

Jarvik, JONATHAN W., FREDERICK D. Reinhart, MICHAEL R. Kuchka, and SALLY A. Adler. 1984. “Altered Flagellar Size-control in Shf-1 Short-flagella Mutants of Chlamydomonas Reinhardtii.” The Journal of Protozoology 31 (2): 199–204.

Kong, Ji Na, Kara Hardin, Michael Dinkins, Guanghu Wang, Qian He, Tarik Mujadzic, Gu Zhu, Jacek Bielawski, Stefka Spassieva, and Erhard Bieberich. 2015. “Regulation of Chlamydomonas Flagella and Ependymal Cell Motile Cilia by Ceramide-Mediated Translocation of GSK3.” Molecular Biology of the Cell 26 (24): 4451–65. https://doi.org/10.1091/mbc.E15-06-0371.

Laiman, Jessica, Julie Loh, Wei-Chun Tang, Mei-Chun Chuang, Hui-Kang Liu, Bi-Chang Chen, Yi-Cheng Chang, Lee-Ming Chuang, and Ya-Wen Liu. 2021. “Dynamin-2 Phosphorylation as A Critical Regulatory Target of Bin1 and GSK3α for Endocytosis in Muscle.” BioRxiv, January, 2021.10.11.463889. https://doi.org/10.1101/2021.10.11.463889.

Lefebvre, P. A., S. A. Nordstrom, J. E. Moulder, and J. L. Rosenbaum. 1978. “Flagellar Elongation and Shortening in Chlamydomonas. IV. Effects of Flagellar Detachment, Regeneration, and Resorption on the Induction of Flagellar Protein Synthesis.” The Journal of Cell Biology 78 (1): 8–27. https://doi.org/10.1083/jcb.78.1.8.

Lefebvre, Paul A. 1995. “Flagellar Amputation and Regeneration in Chlamydomonas.” In Methods in Cell Biology, 47:3–7. Elsevier.

Levy, Elinor Miller. 1974. “Flagellar Elongation as a Moving Boundary Problem.” Bulletin of Mathematical Biology 36: 265–73.

Li, Xiaobo, Weronika Patena, Friedrich Fauser, Robert E. Jinkerson, Shai Saroussi, Moritz T. Meyer, Nina Ivanova, et al. 2019. “A Genome-Wide Algal Mutant Library and Functional Screen Identifies Genes Required for Eukaryotic Photosynthesis.” Nature Genetics 51 (4): 627–35. https://doi.org/10.1038/s41588-019-0370-6.

Ludington, William B., Hiroaki Ishikawa, Yevgeniy V. Serebrenik, Alex Ritter, Rogelio A. Hernandez-Lopez, Julia Gunzenhauser, Elisa Kannegaard, and Wallace F. Marshall. 2015. “A Systematic Comparison of Mathematical Models for Inherent Measurement of Ciliary Length: How a Cell Can Measure Length and Volume.” Biophysical Journal 108 (6): 1361–79. https://doi.org/10.1016/j.bpj.2014.12.051.

Marshall, Wallace F. 2015. “How Cells Measure Length on Subcellular Scales.” Special Issue: Quantitative Cell Biology 25 (12): 760–68. https://doi.org/10.1016/j.tcb.2015.08.008.

Marshall, Wallace F, Hongmin Qin, Mónica Rodrigo Brenni, and Joel L Rosenbaum. 2005. “Flagellar Length Control System: Testing a Simple Model Based on Intraflagellar Transport and Turnover.” Molecular Biology of the Cell 16 (1): 270–78. https://doi.org/10.1091/mbc.e04-07-0586.

Marshall, Wallace F., and Joel L. Rosenbaum. 2001. “Intraflagellar Transport Balances Continuous Turnover of Outer Doublet Microtubules: Implications for Flagellar Length Control.” The Journal of Cell Biology 155 (3): 405–14.

Miki, Daisuke, Yuki Kobayashi, Tomoya Okada, Tatuso Miyamoto, Nobuyuki Takei, Yuko Sekino, Noriko Koganezawa, Tomoaki Shirao, and Yumiko Saito. 2019. “Characterization of Functional Primary Cilia in Human Induced Pluripotent Stem Cell-Derived Neurons.” Neurochemical Research 44 (7): 1736–44. https://doi.org/10.1007/s11064-019-02806-4.

Miyoshi, Ko, Kyosuke Kasahara, Ikuko Miyazaki, and Masato Asanuma. 2009. “Lithium Treatment Elongates Primary Cilia in the Mouse Brain and in Cultured Cells.” Biochemical and Biophysical Research Communications 388 (4): 757–62. https://doi.org/10.1016/j.bbrc.2009.08.099.

Nachury, Maxence V., E. Scott Seeley, and Hua Jin. 2010. “Trafficking to the Ciliary Membrane: How to Get across the Periciliary Diffusion Barrier?” Annual Review of Cell and Developmental Biology 26: 59–87. https://doi.org/10.1146/annurev.cellbio.042308.113337.

Nakamura, Shogo, Hiroyoshi Takino, and Manabu K. Kojima. 1987. “Effect of Lithium on Flagellar Length in Chlamydomonas Reinhardtii.” Cell Structure and Function 12 (4): 369–74. https://doi.org/10.1247/csf.12.369.

Nguyen, Rachel L., Lai-Wa Tam, and Paul A. Lefebvre. 2005. “The LF1 Gene of Chlamydomonas Reinhardtii Encodes a Novel Protein Required for Flagellar Length Control.” Genetics 169 (3): 1415–24. https://doi.org/10.1534/genetics.104.027615.

Onishi, Masayuki, James G. Umen, Frederick R. Cross, and John R. Pringle. 2019. “Cleavage-Furrow Formation without F-Actin in <em>Chlamydomonas</Em>.” BioRxiv, January, 789016. https://doi.org/10.1101/789016.

Osterman, Natan, and Andrej Vilfan. 2011. “Finding the Ciliary Beating Pattern with Optimal Efficiency.” Proceedings of the National Academy of Sciences 108 (38): 15727–32.

Ou, Young, Camila Dores, Jose-Rafael Rodriguez-Sosa, Frans A. van der Hoorn, and Ina Dobrinski. 2014. “Primary Cilia in the Developing Pig Testis.” Cell and Tissue Research 358 (2): 597–605. https://doi.org/10.1007/s00441-014-1973-y.

Ou, Young, Yibing Ruan, Min Cheng, Joanna J Moser, Jerome B Rattner, and Frans A van der Hoorn. 2009. “Adenylate Cyclase Regulates Elongation of Mammalian Primary Cilia.” Experimental Cell Research 315 (16): 2802–17. https://doi.org/10.1016/j.yexcr.2009.06.028.

Ou, Young, Ying Zhang, Min Cheng, Jerome B. Rattner, Ina Dobrinski, and Frans A. van der Hoorn. 2012. “Targeting of CRMP-2 to the Primary Cilium Is Modulated by GSK-3β.” PloS One 7 (11): e48773. https://doi.org/10.1371/journal.pone.0048773.

Reiter, Jeremy F., and Michel R. Leroux. 2017. “Genes and Molecular Pathways Underpinning Ciliopathies.” Nature Reviews Molecular Cell Biology 18 (9): 533–47. https://doi.org/10.1038/nrm.2017.60.

Rohatgi, Rajat, and William J Snell. 2010. “The Ciliary Membrane.” Current Opinion in Cell Biology 22 (4): 541–46. https://doi.org/10.1016/j.ceb.2010.03.010.

Rosenbaum, J. L., J. E. Moulder, and D. L. Ringo. 1969. “Flagellar Elongation and Shortening in Chlamydomonas. The Use of Cycloheximide and Colchicine to Study the Synthesis and Assembly of Flagellar Proteins.” The Journal of Cell Biology 41 (2): 600–619. https://doi.org/10.1083/jcb.41.2.600.

Smillie, Karen J., and Michael A. Cousin. 2012. “Akt/PKB Controls the Activity-Dependent Bulk Endocytosis of Synaptic Vesicles.” Traffic (Copenhagen, Denmark) 13 (7): 1004–11. https://doi.org/10.1111/j.1600-0854.2012.01365.x.

Soave, Arianna, Loraine L. Y. Chiu, Aisha Momin, and Stephen D. Waldman. 2022. “Lithium Chloride-Induced Primary Cilia Recovery Enhances Biosynthetic Response of Chondrocytes to Mechanical Stimulation.” Biomechanics and Modeling in Mechanobiology, January. https://doi.org/10.1007/s10237-021-01551-4.

Srinivasan, Saipraveen, Christoph J. Burckhardt, Madhura Bhave, Zhiming Chen, Ping-Hung Chen, Xinxin Wang, Gaudenz Danuser, and Sandra L. Schmid. 2018. “A Noncanonical Role for Dynamin-1 in Regulating Early Stages of Clathrin-Mediated Endocytosis in Non-Neuronal Cells.” PLOS Biology 16 (4): e2005377. https://doi.org/10.1371/journal.pbio.2005377.

Tam, Daniel, and AE Hosoi. 2011. “Optimal Feeding and Swimming Gaits of Biflagellated Organisms.” Proceedings of the National Academy of Sciences 108 (3): 1001–6.

Tam, Lai-Wa, Nedra F. Wilson, and Paul A. Lefebvre. 2007. “A CDK-Related Kinase Regulates the Length and Assembly of Flagella in Chlamydomonas.” The Journal of Cell Biology 176 (6): 819–29. https://doi.org/10.1083/jcb.200610022.

Thompson, Clare L., Anna Wiles, C. Anthony Poole, and Martin M. Knight. 2016. “Lithium Chloride Modulates Chondrocyte Primary Cilia and Inhibits Hedgehog Signaling.” FASEB Journal: Official Publication of the Federation of American Societies for Experimental Biology 30 (2): 716–26. https://doi.org/10.1096/fj.15-274944.

Tinevez, Jean-Yves, Nick Perry, Johannes Schindelin, Genevieve M. Hoopes, Gregory D. Reynolds, Emmanuel Laplantine, Sebastian Y. Bednarek, Spencer L. Shorte, and Kevin W. Eliceiri. 2017. “TrackMate: An Open and Extensible Platform for Single-Particle Tracking.” Image Processing for Biologists 115 (February): 80–90. https://doi.org/10.1016/j.ymeth.2016.09.016.

Wilson, Nedra F, and Paul A Lefebvre. 2004. “Regulation of Flagellar Assembly by Glycogen Synthase Kinase 3 in Chlamydomonas Reinhardtii.” Eukaryotic Cell 3 (5): 1307–19. https://doi.org/10.1128/EC.3.5.1307-1319.2004.

