## Supplemental Material for "Lithium-induced ciliary lengthening sparks Arp2/3 complex-dependent endocytosis"

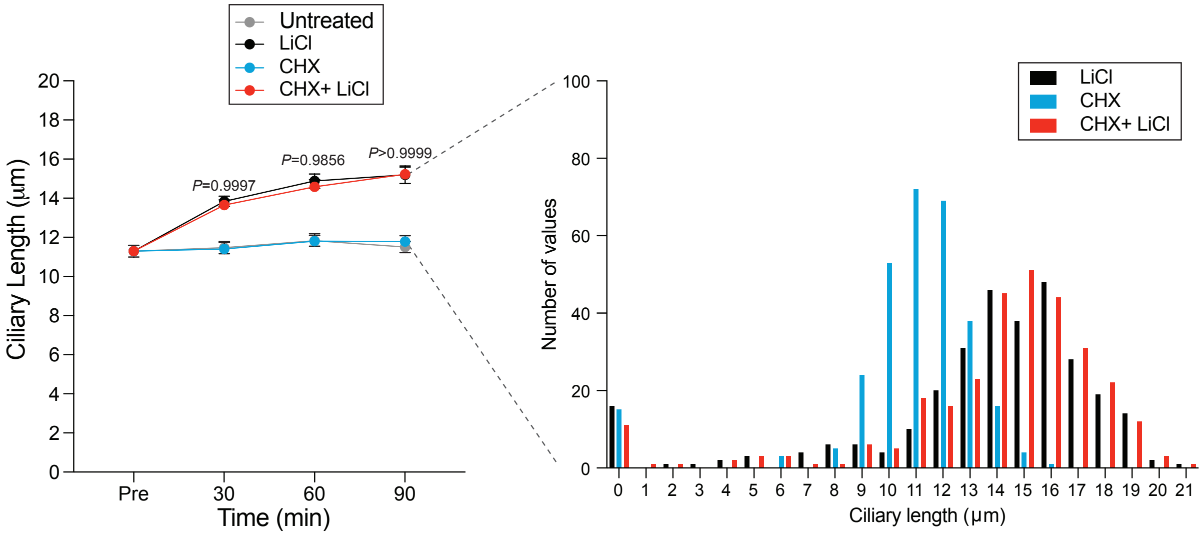


**Supplemental Figure 1. New protein synthesis is not required for lithium-induced ciliary elongation. A)** Wild-type cells were treated with either 25 mM LiCl, 10 µM Cycloheximide (CHX), or a combination of the two drugs. n=30 for each sample at each time point in 3 separate biological replicates. Significance was determined by one-way ANOVA and a Tukey's multiple comparisons test. The p values above the lines show the comparison between cells treated with LiCl alone or LiCl and CHX. In all cases, the comparison is not significant. **B)** A histogram representation of the 90-minute time point from (A). For this, n=100 for each sample in 3 separate biological replicates (300 total points).

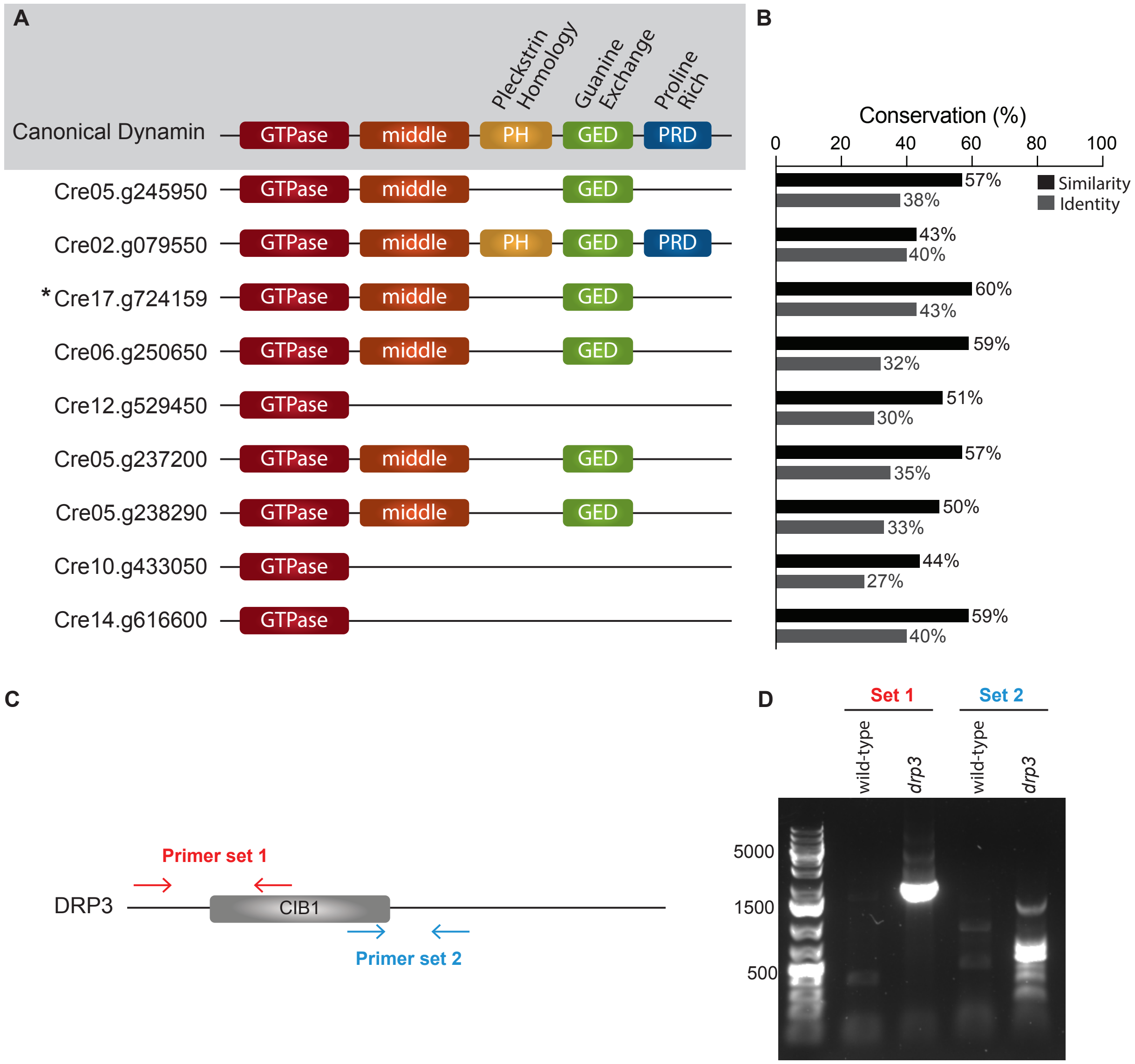


**Supplemental Figure 2. *Chlamydomonas* contains several dynamin related proteins (DRPs) with the most similar to canonical dynamin being DRP3 (Cre17.g724159) A)** The canonical dynamin contains 5 domains represented in different colors above. The *Chlamydomonas* genome contains no conventional dynamins but does contain 9 dynamin related proteins (DRPs) that have various similarities compared with canonical dynamin. The *Chlamydomonas* DRPs above are represented by the gene identifier and are ordered by DRP number (the first is DRP1, the second is DRP2, etc). **B)** The graph on the right shows the similarity and identity of each DRP compared to canonical mammalian dynamin as determined by MUSCLE alignment. The DRP that is the most closely related to canonical dynamin is DRP3, Cre17.g724159. **C)** Schematic of the DRP3 genomic sequence with the primers used to confirm mutation represented by arrows. **D)** DNA gel electrophoresis showing resulting DNA sequences from the PCR using the primer sets shown in C.
